# Validation of two-dimensional video-based inference of finger kinematics with pose estimation

**DOI:** 10.1101/2022.06.22.497125

**Authors:** Letizia Gionfrida, Wan M. R. Rusli, Anil Antony Bharath, Angela E. Kedgley

## Abstract

Accurate capture finger of movements for biomechanical assessments has typically been achieved within laboratory environments through the use of physical markers attached to a participant’s hands. However, such requirements can narrow the broader adoption of movement tracking for kinematic assessment outside these laboratory settings, such as in the home. Thus, there is the need for markerless hand motion capture techniques that are easy to use and accurate enough to evaluate the complex movements of the human hand. Several recent studies have validated lower-limb kinematics obtained with a marker-free technique, OpenPose. This investigation examines the accuracy of OpenPose, when applied to images from single RGB cameras, against a ‘gold standard’ marker-based optical motion capture system that is commonly used for hand kinematics estimation. Participants completed four single-handed activities with right and left hands, including hand abduction and adduction, radial walking, metacarpophalangeal (MCP) joint flexion, and thumb opposition. Accuracy of finger kinematics was assessed using the root mean square error. Mean total active flexion was compared using the Bland–Altman approach, and coefficient of determination of a linear regression. Results showed good agreement for abduction and adduction and thumb opposition activities. Lower agreement between the two methods was observed for radial walking (mean difference between the methods of 5.03°) and MCP flexion (mean difference of 6.82°) activities, due to occlusion. This investigation demonstrated that OpenPose, applied to videos captured with monocular cameras, can be used for markerless motion capture for finger tracking with an error below than 11° and on the order of that which is accepted clinically.

**Author summary:** Decreased hand mobility may limit functionality, and its quantification is fundamental to assess underlying impairments. Optical motion capture technologies are the most accurate means by which to quantify hand motion. As this approach involves placing markers on the skin and recording hand movements using multiple cameras, there are limitations of physical space, time requirements, and financial implications. Therefore, the adoption of these practices is confined to laboratory settings. In clinical settings, goniometry is used to quantify hand range of motion (ROM), but this also involves lengthy processes and requires face-to-face assessments. Alternative solutions have been investigated to quantify hand mobility remotely and support home-based care interventions. However, none has been shown to be accurate enough to replace the gold-standard measurement of hand ROM in clinical settings. Recently, markerless technologies that leverage artificial intelligence have exhibited great potential for human movement analysis, but these studies have validated markerless tracking technologies for the lower limb only. We demonstrate that the validity of these models can be extended to capture hand mobility, making it also possible to assess hand function remotely.

## Introduction

Optical marker-based motion tracking technologies can be classified based upon their working principle, dividing them into marker-based and markerless [1]. Marker-based motion capture relies on either active infrared (IR) or passive retroreflective markers whose motion is tracked by two or more cameras Passive optical marker-based settings are considered the ‘gold standard’ to measure kinematics in the field of hand biomechanics [2]. However, conventional marker-based motion capture systems are expensive, confined to the laboratory, not easily accessible to the broad population, and time consuming to set up, thus are difficult to adopt in clinical settings [3, 4].

Advances in machine learning have allowed computer vision researchers to gather fully labelled images and train neural networks to automatically detect the positions of users’ anatomical landmarks from video. Recently, several computational tools have emerged as potential platforms for 2D markerless tracking and pose estimation, such as OpenPose [5] or DeepLabCut [6]. However, while the hand biomechanics community demands accuracies on the order of 1°, and established instrument error of clinical universal goniometers is 6.6° [7], the validity of markerless tracking is usually outside the reach of clinical biomechanics research. Indeed Seethapathi et al. suggested that the implementation of deep-learning-based pose tracking have not prioritized features that matter for movement biomechanics, and the question on whether these models could be extended to clinical biomechanics remains open [8].

Nakano et al. [9] quantified the accuracy of lower limb movements from video data captured using multiple RGB cameras against a markered optoelectronic motion capture technology. They used a direct linear transformation [10] to estimate 3D coordinates of shoulder, elbow, wrist, hip, knee, and ankle joints, from the 2D anatomical landmarks (keypoints) obtained using OpenPose, showing an inaccuracy of 3 cm. Openpose processed the location and then they measured the kinematics post-hoc from the OpenPose outputs. An improved approach was presented by D’Antonio et al. [11], who implemented a pipeline consisting of two RGB cameras and a linear triangulation algorithm to convert 2D coordinates obtained with OpenPose in a 3D coordinate system. Results showed that their system could track lower limb segment angles relative to the global frame with errors of up to 9.9°. However, the choice to use two cameras may restrict the utilization of videos recorded in the home or other common settings.

Most recently, OpenPose has been assessed for markerless motion capture of gait using a single camera. Sakurai et al. [12] compared 3D gait kinematics acquired with a markered optoelectronic motion capture system against 2D keypoints extracted from a single video camera. Their study presented an error of approximately 5° between the systems. Similarly, Stenum et al. [13] compared 2D sagittal gait kinematics estimated using OpenPose against 3D motion capture, showing errors in flexion-extension of 4.0° for the hip, 5.6° for the knee, and 7.4° for the ankle. Finally, Drazan et al. [14] assessed the performance of OpenPose against a marker-based motion capture system in estimating lower limb angles in the 2D sagittal plane during vertical jump. They obtained errors lower than 3.22° in flexion-extension across the hip, knee, and ankle when the two methods were compared. However, it remains unclear whether the validation of OpenPose can be extended to address the specific needs of finger kinematics.

The objective assessment of finger kinematics is fundamental to enhance the knowledge of hand functionality in both healthy and impaired populations. Therefore, this work aims to compare 3D kinematics obtained with a gold standard marker-based optoelectronic motion capture system against 2D coronal hand kinematics obtained from a monocular RGB camera using OpenPose. The 3D motion representations were automatically projected on the 2D image frames captured using a synchronized video camera to compare 3D kinematics in 2D.

## Results

We collected a comprehensive dataset of hand images containing 3D motion capture parameters obtained with from an optoelectronic motion capture system and coronal plane video recordings of healthy adults. The pipeline implemented in investigation is in Fig 1. The 3D keypoints (anatomical landmarks) were filtered and mapped onto the two-dimensional video recordings to extract ground truth kinematics. The RGB frames were processed using OpenPose, subsequently filtered, and the hand kinematics were extracted and compared against the ground truth kinematics.

**Fig 1.**
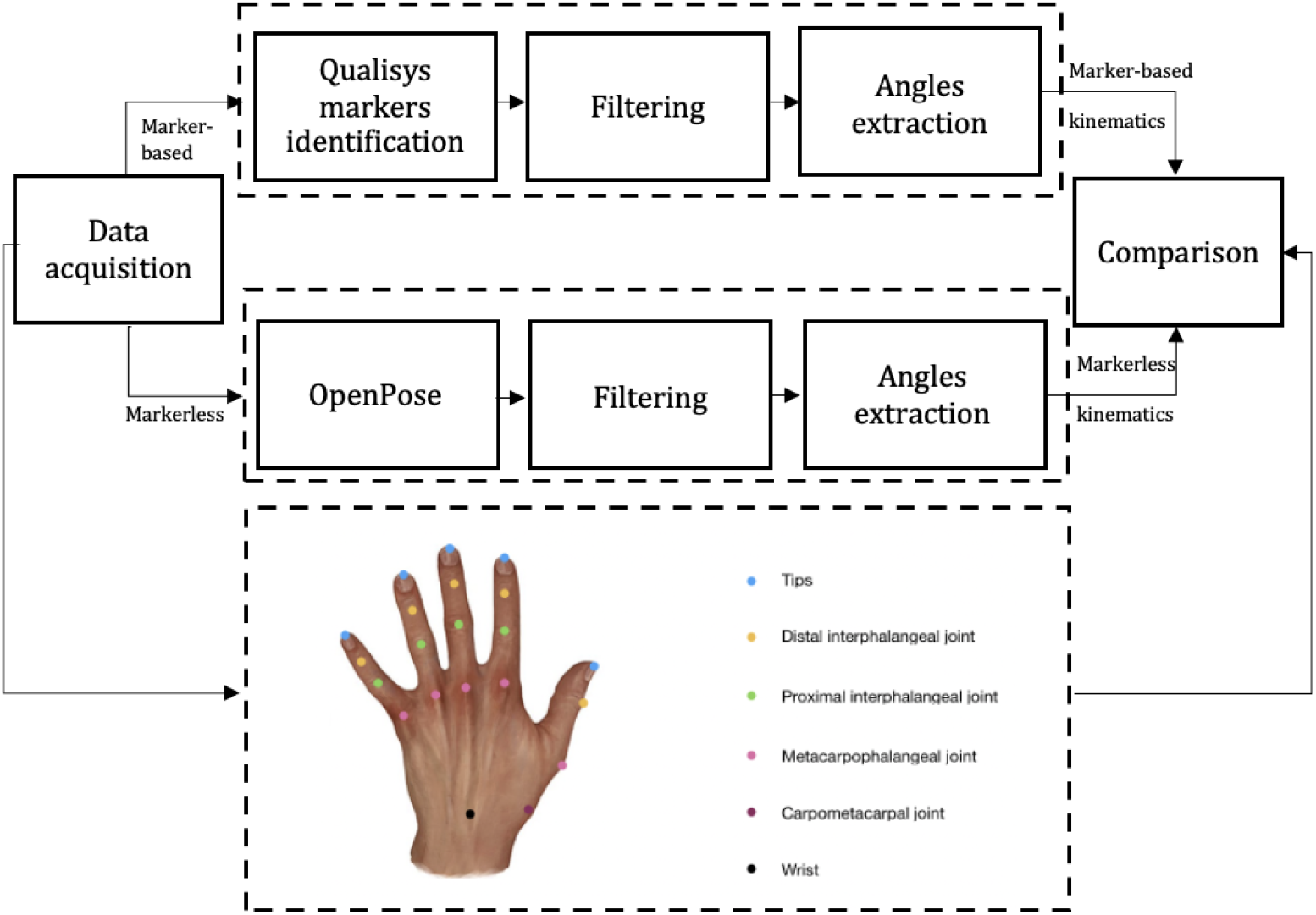
Overview of the workflow. Flowchart of the experimental set-up of the marker-based versus markerless investigation.

Representative plots for abduction and adduction, radial walking, metacarpophalangeal flexion and thumb opposition in Fig. 2 show the similarity between the two trends determined using OpenPose and obtained with the optoelectronic motion capture system, during the four tasks performed.

**Fig 2.**
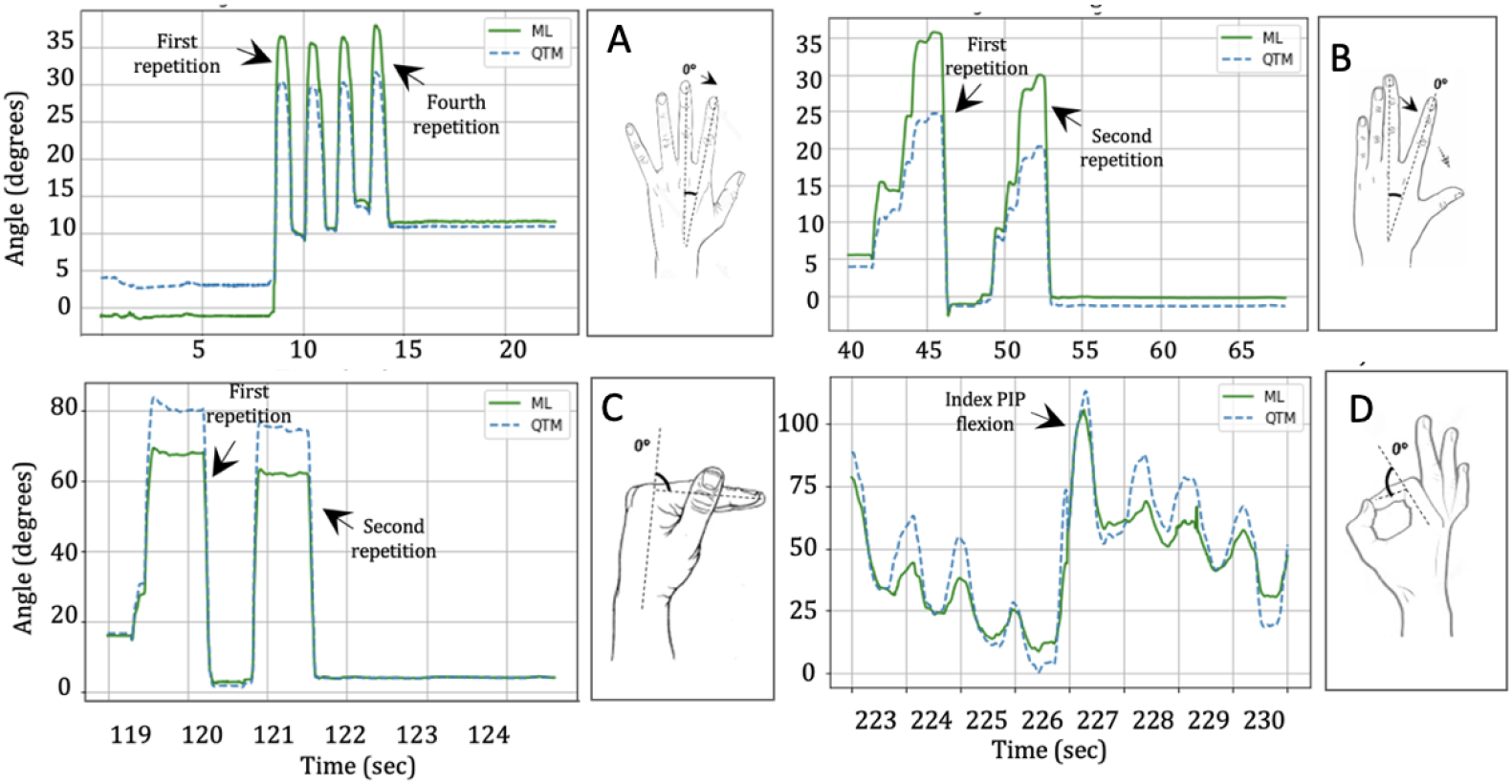
Examples of raw data for one healthy participant. Examples of averaged raw data for (A) 2nd-to-3rd digit angle for four repetitions of the abduction and adduction task, (B) 2nd-to-3rd finger angle for two repetitions of the radial walking task, (C) 2nd metacarpophalangeal (MCP) joint angle for two repetitions of the MCP flexion task, (D) 2nd proximal interphalangeal joint angle for the thumb opposition task, estimated using OpenPose (ML; solid lines) and measured with the optoelectronic system (QTM; dashed lines) for one representative healthy participant.

As a metric of comparison of the two-time series, once the angles were obtained from the two tracking techniques, the differences were computed using the root mean square error (RMSE).The total active flexion (TAF) was extracted for each digit and for each of the exercises under inspection, Bland–Altman plots and linear regression were used to illustrate the agreement between the methodologies. In Bland–Altman analysis the agreement between two measures is assessed with the estimation of the standard deviation (SD) of differences with 95% limits of agreement (LoA) ± 1.96 SDs of the mean.

For abduction and adduction, the finger kinematics inferred with OpenPose presented an RMSE below 9° (Fig 3),with larger errors observed for the 4th-to-5th digit angle due to occlusion by the other fingers while performing the task. The total active flexion (TAF) values exhibited a mean difference between OpenPose and the optoelectronic motion capture system of 4.72° (Fig 4) with limits of agreement (LoA) of 8.8° and 0.56°, and coefficient of determination of 0.73, indicating good agreement (reference) between the two methodologies for this activity.

**Fig 3.**
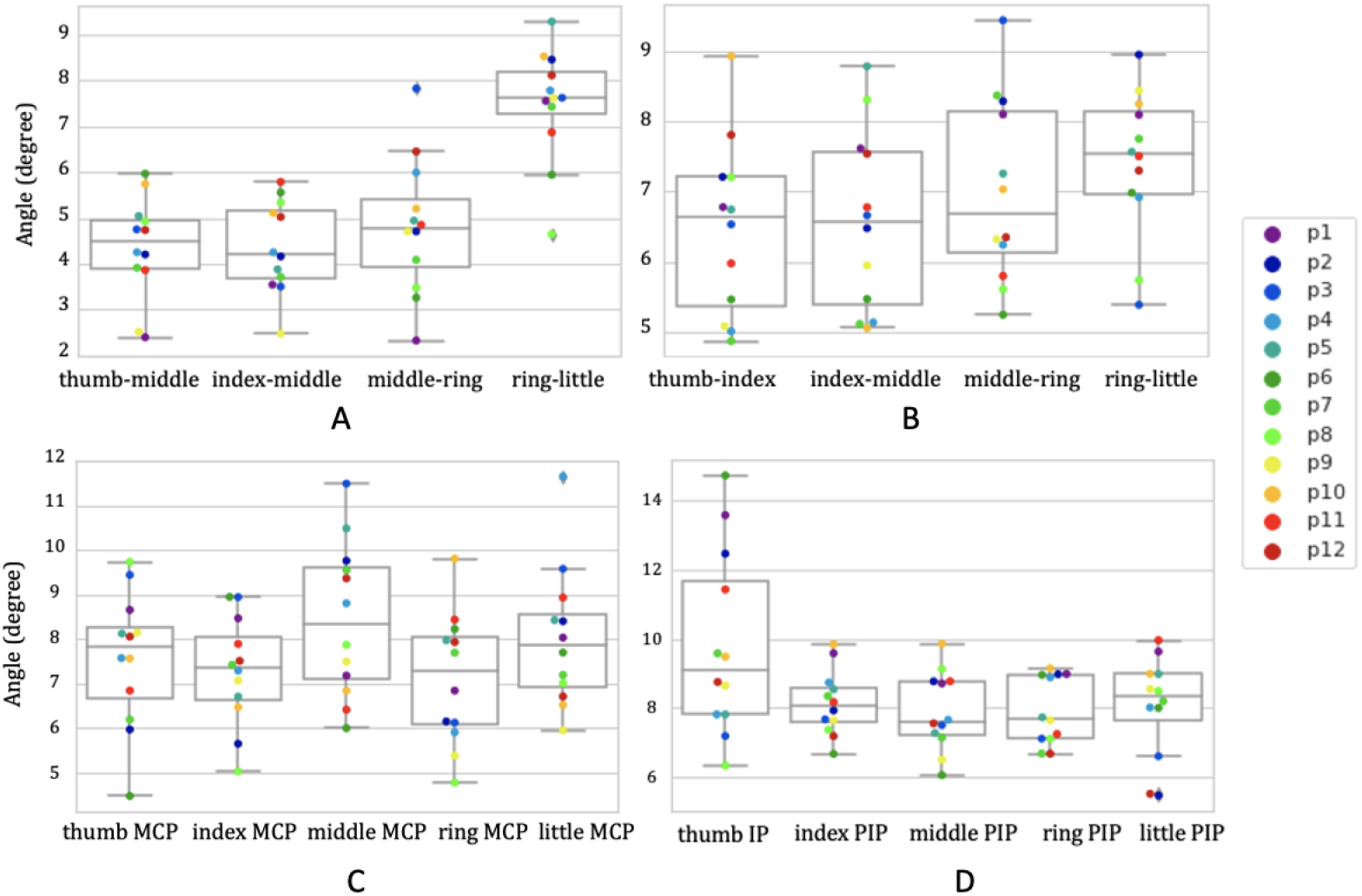
RMSEs for the four activities. Root mean square error (RMSE) differences between OpenPose on monocular images and marker-based optoelectronic system during A) finger abduction and adduction, B) radial walking, C) finger metacarpophalangeal flexion, and D) thumb opposition. Each colour represents a different participant.

**Fig 4.**
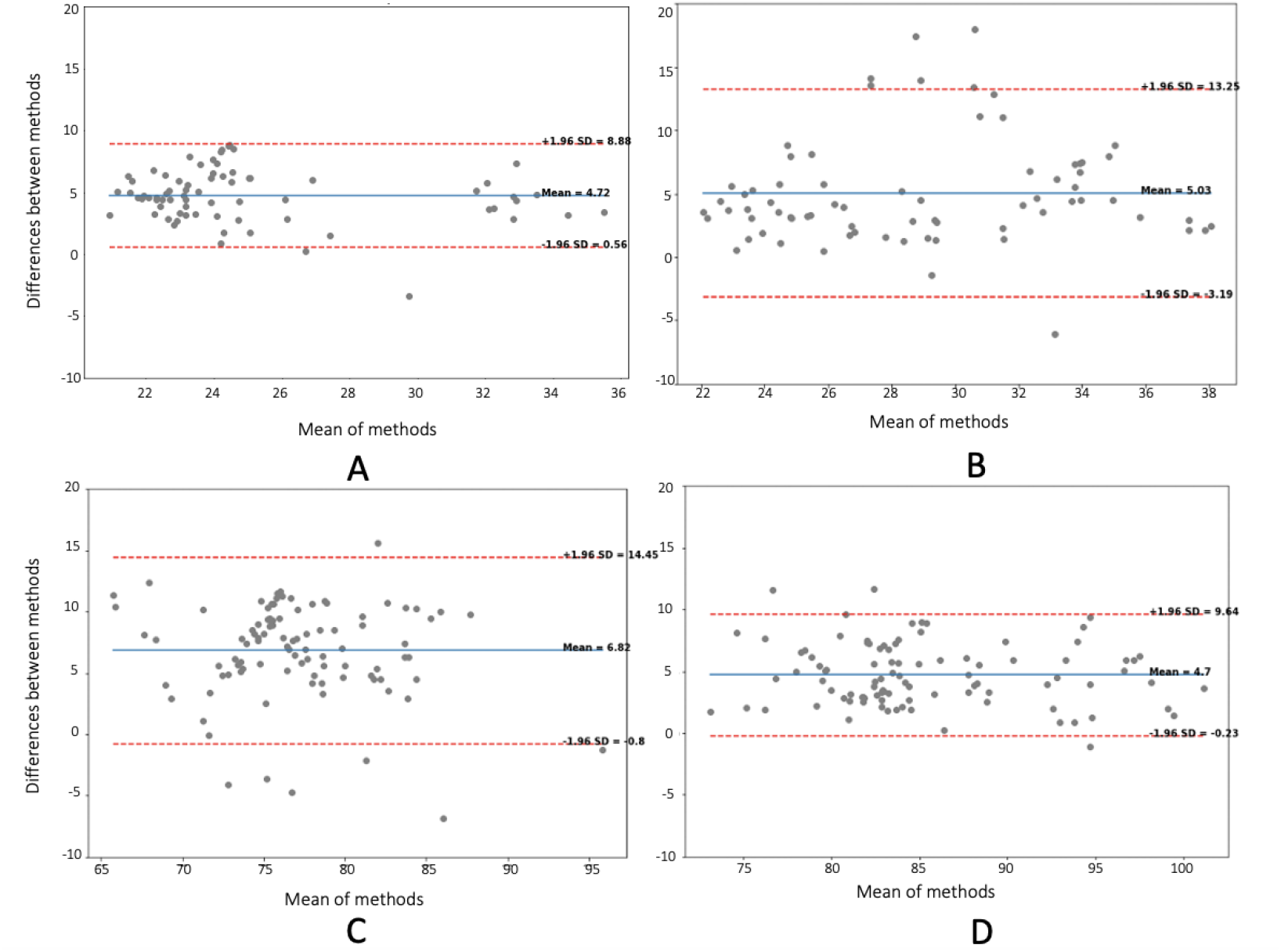
Bland-Altman. Bland-Altman plots plots of total active flexion for A) abduction and adduction, B) radial walking, C) metacarpophalangeal flexion, and D) thumb opposition of the 2nd, 3rd, 4th and 5th digits for the metacarpophalangeal and proximal interphalangeal joints of the left and the right hands.

For the radial walking hand activity performed on the table, the finger kinematics estimated with OpenPose presented an RMSE below 9° (Fig 3). The TAF values (fig 6B) presented a mean difference between the methods of 5.03° with LoA ranging from 13.25° to -3.19° (Fig 4). Larger variability (coefficient of determination=0.40) was suggested, as compared to the abduction and adduction activity.

**Fig 5.**
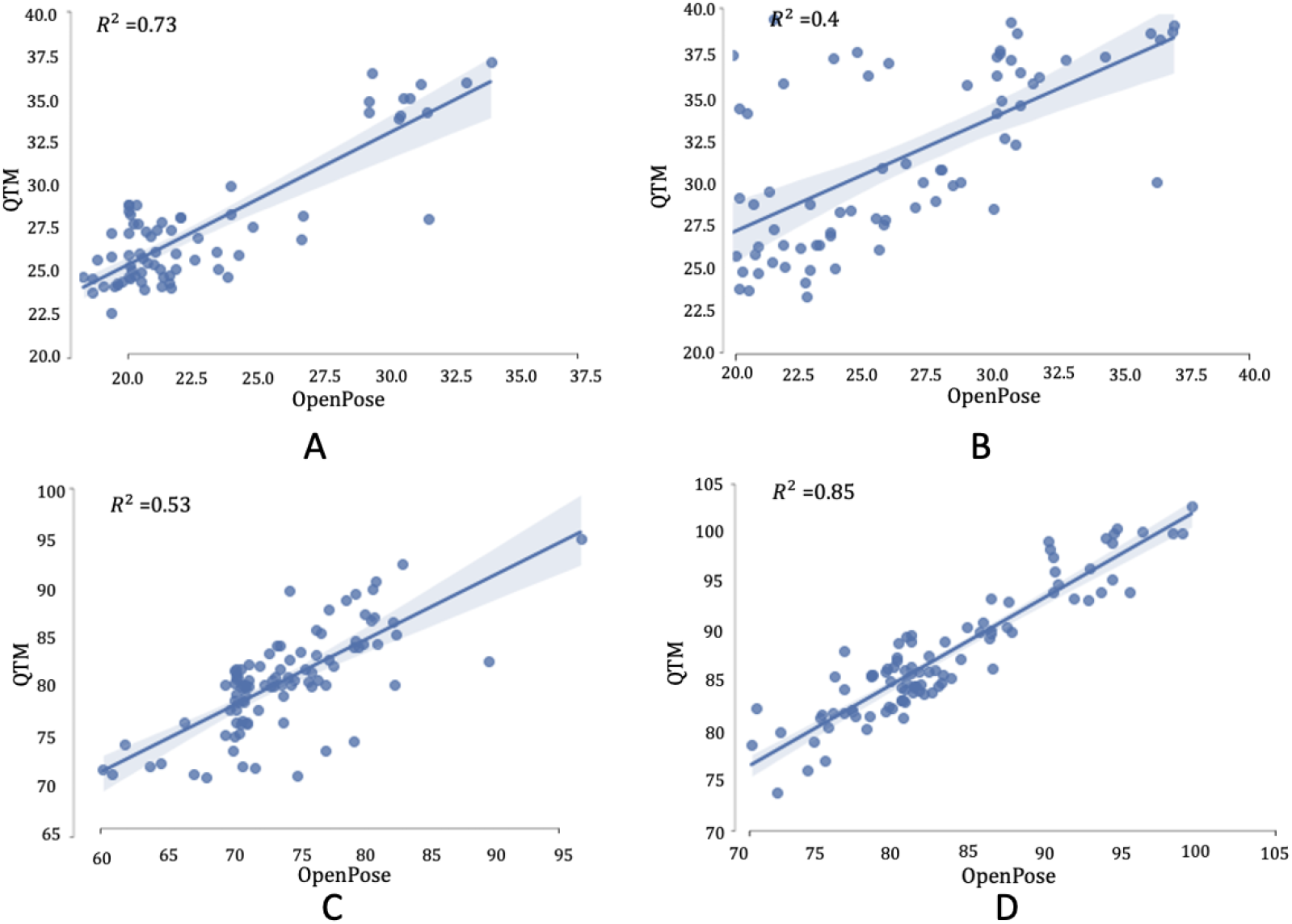
Linear regression. Linear regression plots of total active flexion for (A) abduction and adduction, (B) radial walking, (C) MCP flexion and (D) thumb opposition of the 2nd, 3rd, 4th and 5th digits for the metacarpophalangeal and proximal interphalangeal joints of the left and the right hands.

**Fig 6.**
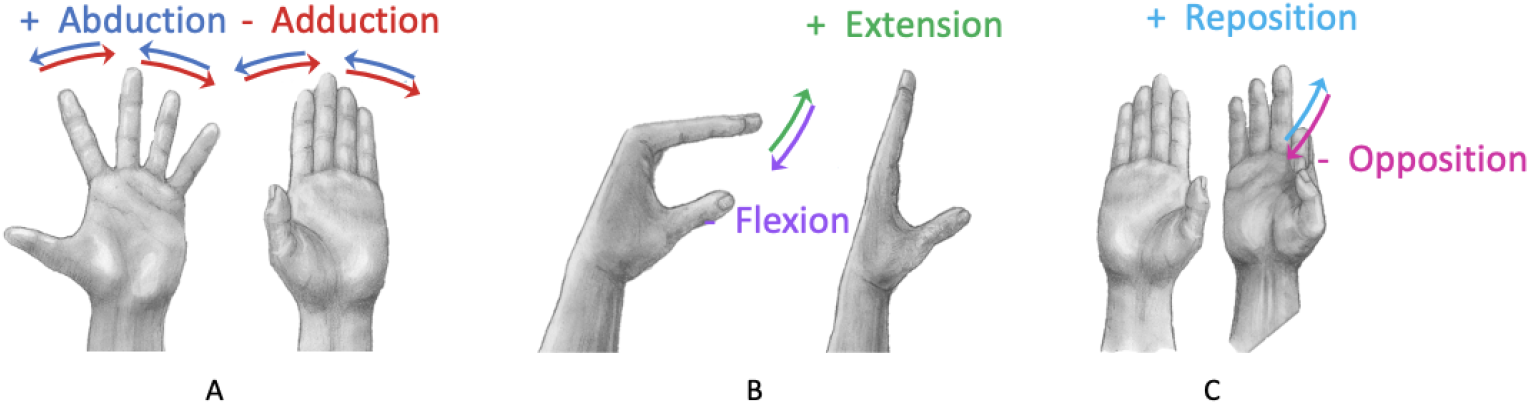
Different hand exercises. Illustrating: A) abduction and adduction, B) metacarpophalangeal flexion, and C) thumb opposition.

During the metacarpophalangeal joint flexion activity the comparison between the two methodologies presented an error below 11° Fig 3), apart from two participants who had an error value between 11° and 12°. The Bland-Altman plot (Fig 4) presented a mean difference of 6.82° (Fig 4) with LoA that went from 14.45° for the upper limit (+1.96 SD) to -0.8° for the lower limit. The comparison between the two methodologies yielded a modest coefficient of determination value of 0.53.

Finally, during thumb opposition task, the RMSEs (Fig 3) were below 10° for 93.3% of the estimated values, while the other 6.7% reported an error between 12° and 14.5°. The principal reason for observing higher errors in 10% of the cases was occlusion by the other fingers, and OpenPose inadvertently swopping finger segment values. The mean difference between values (Fig 4) was 4.7° with LoA 9.64° and -0.23°, and a coefficient of determination of 0.85.

## Discussion

This work proposes the validation of a tracking measurement to quantify hand kinematics during specific hand activities using a monocular RGB camera. The chosen markerless technique makes use of a convolutional-neural-network-based model, known as OpenPose, and two filtering techniques, the Hampel and the Butterworth, to capture, quantify and evaluate finger kinematics from video recordings. The accuracy of OpenPose in tracking 2D finger kinematics was assessed by comparing it with the 2D projections of 3D finger kinematics obtained using a marker-based motion capture system.

Markerless technologies that leverage deep-learning architectures have exhibited great potential for motion tracking, using monocular video cameras. For instance, two-dimensional pose estimation models have been validated for human gait, reporting an error of 5° to 15° [9, 11, 13]. Leveraging these findings, this paper offers a preliminary proof-of-concept investigation showing that hand pose estimation of hand kinematics using OpenPose can reach similar levels of accuracy to assess kinematics during hand-specific exercises. The comparison between the marker-based and the markerless technologies presented an error below 10°, apart from a few outliers; these occurred with a 3.4% frequency rate.

Differences when comparing the two methodologies may be introduced by several factors, including the nature of the video recording. For instance, OpenPose depends on images labelled with keypoints, whereas marker placement relies on the physical location of anatomical landmarks. Another possible cause of outliers could be linked to the comparison of the two-dimensional keypoints and the 3D motion capture parameters. While we calculated the included angle between two vectors from a projection of the 3D landmarks onto a plane, the fingers were still moving in 3D space, leading to potential differences in the angle calculation. A further potential reason for these outliers was self-occlusion.

Across the different hand exercises illustrated, the coefficient of determination presented good agreement between the two methods for the abduction and adduction and the thumb opposition activities. Lower coefficient of determination values, representing lower agreement between the two methods, were observed for the radial walking and the metacarpophalangeal flexion activities. During the radial walking task, it was noted that the hand positioned vertically reduced the amount of keypoints lost, compared to when the hand was seated on the table. This was due to the nature in which OpenPose was trained to infer hand kinematics from monocular RGB cameras. Given the modest agreement of the two tracking systems during the radial walking task, and since the abduction adduction activity was able to extract the same joint ranges of motion as the radial walking exercise, it is noted that the abduction and adduction task would be the preferred activity for translation into clinical practice using applications monitored using OpenPose. The modest the coefficient of determination value (0.53) observed during the metacarpophalangeal flexion task can be attributed to the fact that during RGB video acquisition the 2nd 3rd, and 4th digits were partially occluded by the 5th digit. Furthermore, it was visually observed that during occlusion OpenPose inverted the tracking, swopping the digits’ values and causing visible errors for 18% of the dataset. This error could be mitigated by adopting visual manual postprocessing techniques or occlusion detection networks. However, this approach could not be automated and thus would limit the adoption of any activity into clinical practice.

OpenPose provides the joint centre locations together with the confidence values. When the confidence value was low, then error unrelated to occlusion, angle calculation, and the nature of the video recording was attributed to intrinsic parameters, as this tracking methodology does not estimate hand movements perfectly from frame-to-frame. The Bland-Altman plots (Fig. 4) illustrated that the biases (mean differences) across the methods were consistent, ranging from 4.7° to 6.8°. Therefore, by offsetting the results with the consistent biases detected in these acquisitions, the accuracy of future results could potentially be improved. Given the constituency of the biases produced in output, further adoption of these findings would include an automated bias-correcting solution.

This investigation has limitations, including the lack of tests under different visualization parameters and lightening conditions and the intrinsic inaccuracy of the tracking system (OpenPose). Also, the selected pre-trained network was chosen as previous studies had validated this model for lower limb kinematics. However, a pre-trained model was utilized, and this model was not trained for the specific hand exercises included in the study.

Another limitation was identified by the extraction of two-dimensional hand keypoints, while the selected architecture (OpenPose) is also able to provide 3D parameters when more than one camera is utilised. The difference in two-dimensional and 3D parameters, as well as discrepancies in capturing the data from using different viewpoints or perspectives (e.g., sagittal, transverse).

The entire approach provides a fully labelled dataset gathered using one monocular camera (e.g., in smartphones/laptops) and encourages researchers to download and our large-scale publicly available dataset and train novel architectures to improve the accuracy of monocular 2D tracking. Given the latest advantages of novel smartphone devices delivered with dual cameras, future investigations could include capturing image from additional cameras, enlarging the capabilities of this current investigation.

Future directions for the research include the evaluation of the selected markerless architecture to impaired hands. In clinical hand biomechanics, hand kinematics are a crucial metric to quantify changes due to degenerative pathologies. This approach could not only be used to monitor patient’s diseases in their natural environments, but also to support remote rehabilitative pathway, supporting objectivity in remote hand therapy and leading to possible improved clinical outcomes and better disease management.

Despite the promising features demonstrated by pose estimation models to track fine movements of human hands, video-annotation and manual segmentation still limits the scalability of this approach to fully automated clinical applications. An approach that would enable automated segmentation and video segment classification, leveraging video-level label data, could extend the capabilities of this investigation into clinical settings and provide the ability to examine larger volumes of video data in uncontrolled environments.

## Materials and methods

### Experimental setup

Twelve healthy volunteers (eight female, four male) participated in the experiment. Participants were asked to attend a single session of recording of hand kinematics. Written informed consent was obtained from each participant. All participants involved in this investigation were healthy volunteers, with no hand impairment. The protocol was approved by the Imperial College Research Ethics Committee (18IC4673). Upon arrival, participants were briefed on the project, guided through a review of the participant information sheet, asked to sign the consent form, and informed of the set of sequences to perform. Written informed consent was obtained from each participant. Participants were visually supported by a PowerPoint presentation that guided them through the hand exercises to be performed with both the right and left hands.

Participants were visually supported by a PowerPoint presentation that guided them through the hand exercises to be performed with both the right and left hands. These were performed with while seated on a standard height chair with both feet flat on the floor. Participants were asked to perform interventions relevant to improving ROM, selected from amongst the hand exercises previously adopted in hand biomechanics studies. The hand activities performed in this investigation were selected to include different numbers of degrees of freedom. The first activity performed was finger abduction and adduction of the 2nd to 5th digits Fig 6. Participants were asked to spread the fingers away from the long 3rd finger (abduction), and then to bring the fingers back, near the 3rd finger (adduction). This was repeated four times for each hand. The second activity was the radial walking exercise, which consisted of placing the hand on a table and sliding the fingers one at a time towards the 1st digit, which was repeated twice for each finger. The third activity was metacarpophalangeal joint flexion Fig 6, where participants were asked to bend the metacarpophalangeal joints of the 2nd to 5th digits twice. The fourth task was thumb opposition Fig 6, where participants were asked to place the pad of the thumb opposite to the 2nd to 5th digits twice bending the proximal interphalangeal (PIP) joint as much as possible. This activity was repeated twice for each hand.

### Marker-based processing

A total of twenty-six passive reflective hemisphere retro-reflective four millimetre markers were placed at specific positions on the dorsal surface of the right wrist, hand, fingers and thumb in accordance with the Hand & Wrist Kinematics (HAWK) [20] protocol. These semi-spherical markers were placed using double-sided adhesive tape, including the first, second, third, fourth and fifth proximal, intermediate, and distal phalanges. Markers were placed directly over the joint centres and on the fingertips on the distal border of the nail.

The 3D joint coordinates gathered from the markers were captured using an eight-camera Qualisys motion capture system (Oqus 500 + cameras, _¡_0.4 mm error, Qualisys AB, Gothenburg, Sweden) and the Qualisys track manager (QTM) software. RGB video data were recorded using an Oqus RGB camera (Qualisys AB, Gothenburg, Sweden). Both the optical motion capture data and the video data were captured at a 30 Hz frame rate. The QTM system was set to capture continuous recordings for 300 seconds for each hand, one hand at a time.

After the calibration and the synchronization of the system with the RGB external camera, participants were visually supported by a PowerPoint presentation that guided them throughout a set of hand tasks described above. These were performed with both the right and the left hands, while seated on a standard height chair with both feet flat on the floor.

Several steps were carried out before extracting the joint angle computation, including labelling, mapping 2d to 3D, filtering, and segmenting the marker-based data.

Automatic Identification of Markers (AIM) is a function in QTM that automatically identifies and labels the trajectories tracked during a recording. Once a model is created, the connections between the markers are defined by the original model, with new trials that can be added to the model to give it additional examples of distances and angles between markers. Adding new trials to an AIM model will help the software apply it more easily to future test participants. Given this feature offered by QTM, a model was created in accordance with the Hand & Wrist Kinematics (HAWK) [20] marker placement.

The 2D multi-view joint location of 3D motion representations were directly projected onto the 2D image frames capture from the synchronized RGB Oqus camera to compare the to compare 3D kinematics obtained with our gold standard, marker-based optoelectronic motion capture system against the 2D coronal hand kinematics obtained from a monocular RGB camera using OpenPose.

Following the labelling, the mapping, smoothing tool in the trajectory editor of the QTM software was used to reduce spikes and noise in the data output from the motion capture system. A 2nd order Butterworth filter with 5 Hz cut off frequency was selected due to the large frame ranges and presence of high-frequency noise. The filter served as a low-pass filter to attenuate information above the 5 Hz cut-off. Finally, the filtered data were manually segmented to isolate the different exercises for both the right and the left hands.

### Markerless data processing

Video data were captured at 30Hz frame rate using an Oqus RGB (Qualisys AB, Gothenburg, Sweden). OpenPose (version 1.7.0) was run with an NVIDIA Tesla K80 GPU under default settings to extract the keypoints. OpenPose, is a library written in C++ using OpenCV and Caffe that detects 21 keypoints on each of the hands. To capture the hand ROM, the video data were first manually segmented and then OpenPose was executed on each frame of the video Fig 7. Data output from OpenPose were visually observed. Instances where the fingers were incorrectly labelled due to the system swopping one finger with another, were manually labelled, assigning the correct value to the respective finger. Other inconsistencies, for instance, those where the fingers were incorrectly labelled and the tracking was missing due to intrinsic problems with OpenPose, were not manually corrected to minimise the required postprocessing and keep the benchmarked scene as close as possible to uncontrolled capturing settings.

**Fig 7.**
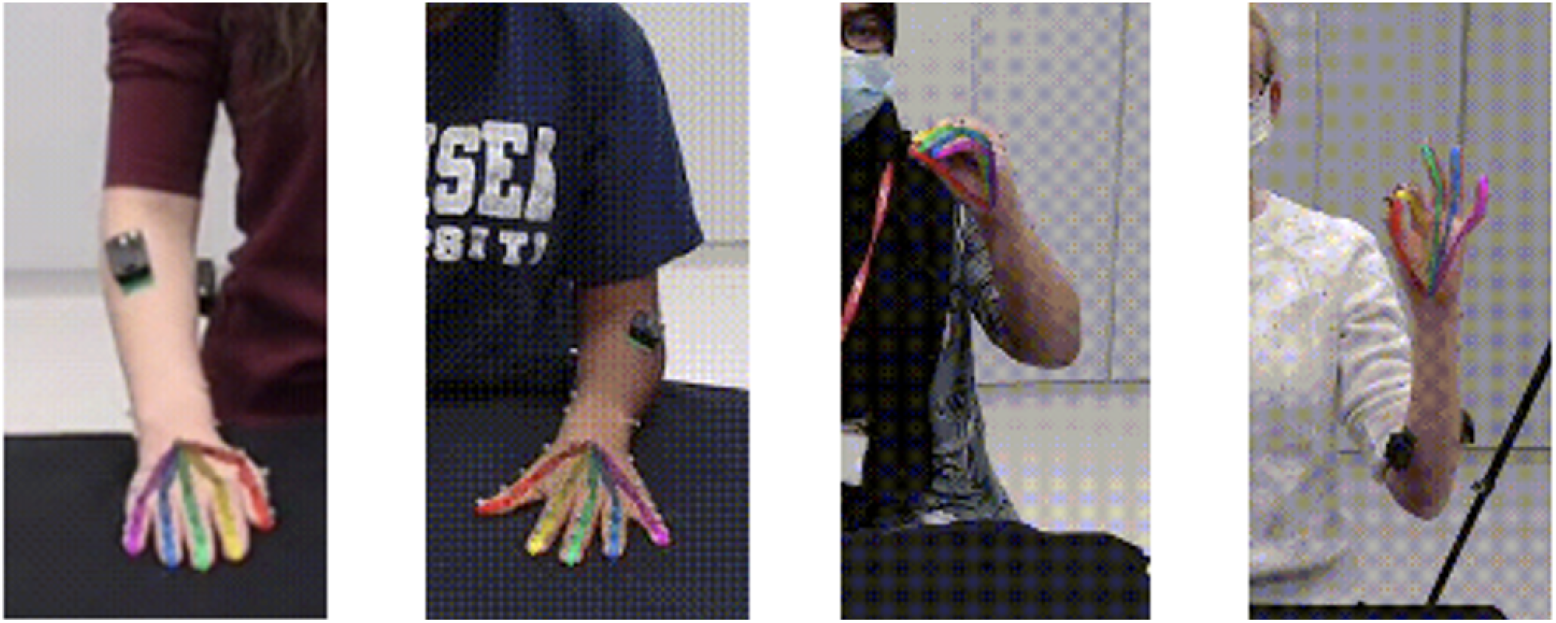
Keypoint visualization. Output from OpenPose that illustrates the inferred keypoints overlapped onto the image frames for four representative participants.

Once the finger keypoints were extracted using OpenPose, four different filtering techniques, previously implemented in similar studies using OpenPose on the lower-limb were tested to prevent the misidentification of keypoints from compromising the ROM detection. The end goal in the evaluation of these filters was i) to select a solution for outliers’ detection, ii) to smooth the raw signal and decrease the noise generated by the architecture.

The filters evaluated were the simple moving average (SMA), Butterworth, and Hampel. To assess the effectiveness of the different approaches, each filter was applied to the thumb opposition sequence of 497 frames (a 16.5-second video with a sampling rate of 30 frames/second). The Hampel filter was the accepted approach for outliers removal. It had two parameters to be tuned, and different configurations were tested (window sized 4, 6, 10, 20 and 60), choosing the multiplying coefficient of the standard deviation (SD) to be kept at one and the window size to be set to four. This setting was found to be able to identify the highest number of visually recognisable outliers when using OpenPose. No threshold was set for what was defined as an “outlier”, opting for a visual inspection of the highest number of outliers identified, as observed in similar lower limb investigations [17].

Following the selection of the multiplying coefficient of the standard deviation and the window size of the Hampel filter for outliers removal, a generalized accepted approach was to smooth the raw signal. Two different filtering techniques were tested, the SMA and the Butterworth. A Butterworth filter with a cut-off frequency of 3 Hz was applied to remove the noise and smooth the signals in output. The cut-off frequency were determined using the residual analysis proposed by Winter et al. [21]. Results of the Butterworth filter for different cut-off frequencies (1 Hz, 2 Hz, and 3 Hz) are illustrated in Fig 8.

**Fig 8.**
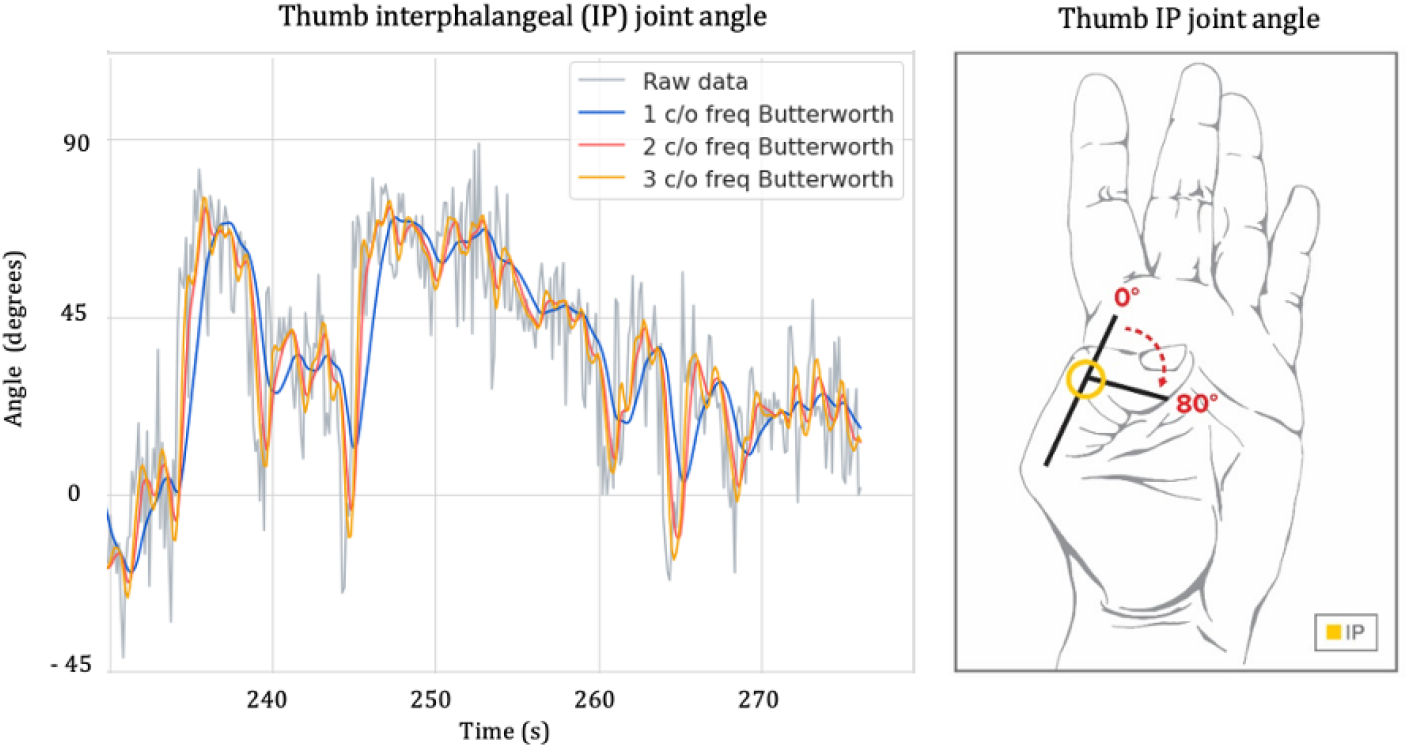
Butterworth filter in output from OpenPose. Butterworth filter with 1 Hz, 2 Hz and 3 Hz cut off frequencies (c/o freq.) applied to the OpenPose signal of the thumb interphalangeal joint angle for one representative participant.

### Hand kinematics

Once the centres of the joints were located using both the maker-based and the markerless motion capture technologies, the hand kinematics were measured. Distal interphalangeal joints were considered to have one degree of freedom (DoF), proximal interphalangeal joints and the two thumbs interphalangeal joints were considered to have one DoF, and metacarpophalangeal joints had two DoF. A total of 36 time-varying angular positions were measured for each participant, with 432 time series extracted for each methodology (marker-based and markerless).

The middle finger was used as a reference for the abduction and adduction task. The eight time-varying angles included the intersection between the thumb and the middle finger (Fig 9), the index and the middle finger, the ring and the middle finger, and little finger and the middle finger, for the left and the right hands. Therefore, eight angles were measured for each participant during the abduction and adduction exercise.

**Fig 9.**
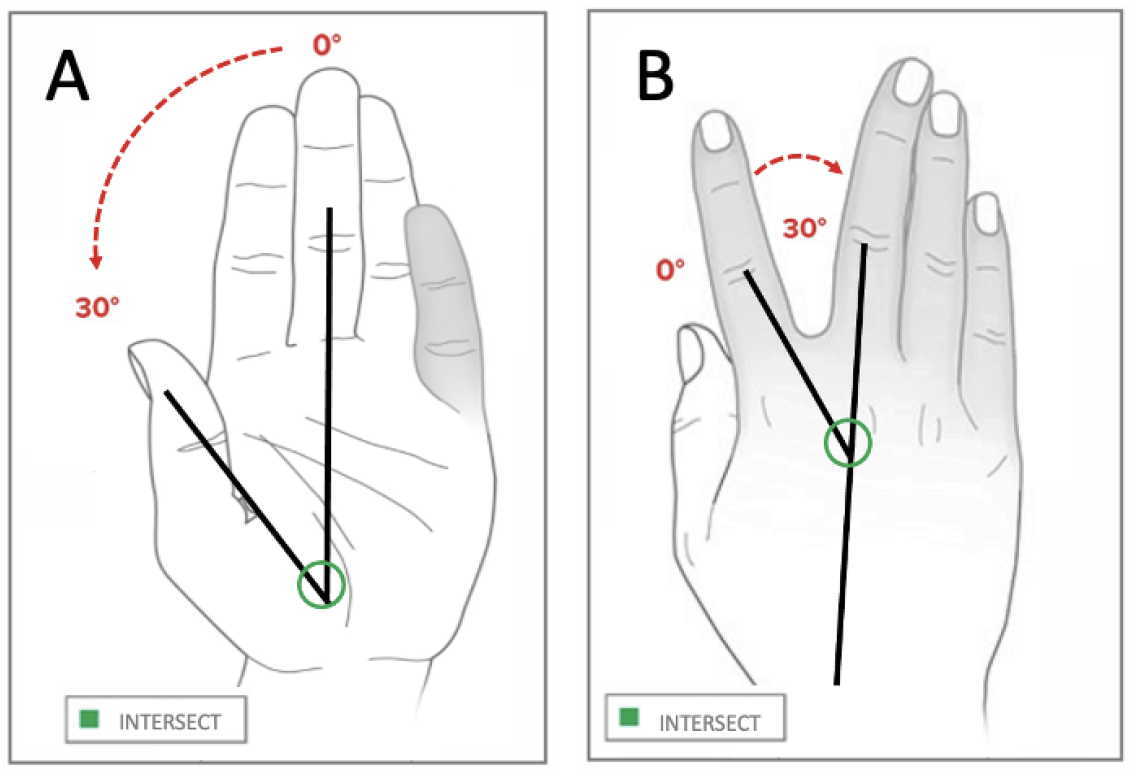
Abduction and addution angles. Measured position for the finger intersect joint of the index finger (A), and of the thumb (B).

During the radial walking task, the reference digit was always considered the one that slid radially prior to digit performing the sliding. The eight angles measured included the intersects between the thumb and the index, the index and the middle (Fig 9), the middle and the ring, and the ring and the little finger, both the right and the left hands.

For the metacarpophalangeal flexion activity the measured angles were the metacarpophalangeal angles of thumb, index, middle, ring, and little fingers for a total of eight angle time series for the right and the left hands (Fig 10). Finally, during the thumb opposition, ten angles were measured. Those angles included the metacarpophalangeal joint angles of the thumb, the interphalangeal joints of the thumbs, and the proximal interphalangeal joints angles of the index, the middle, the ring, and the little finger (Fig 10).

**Fig 10.**
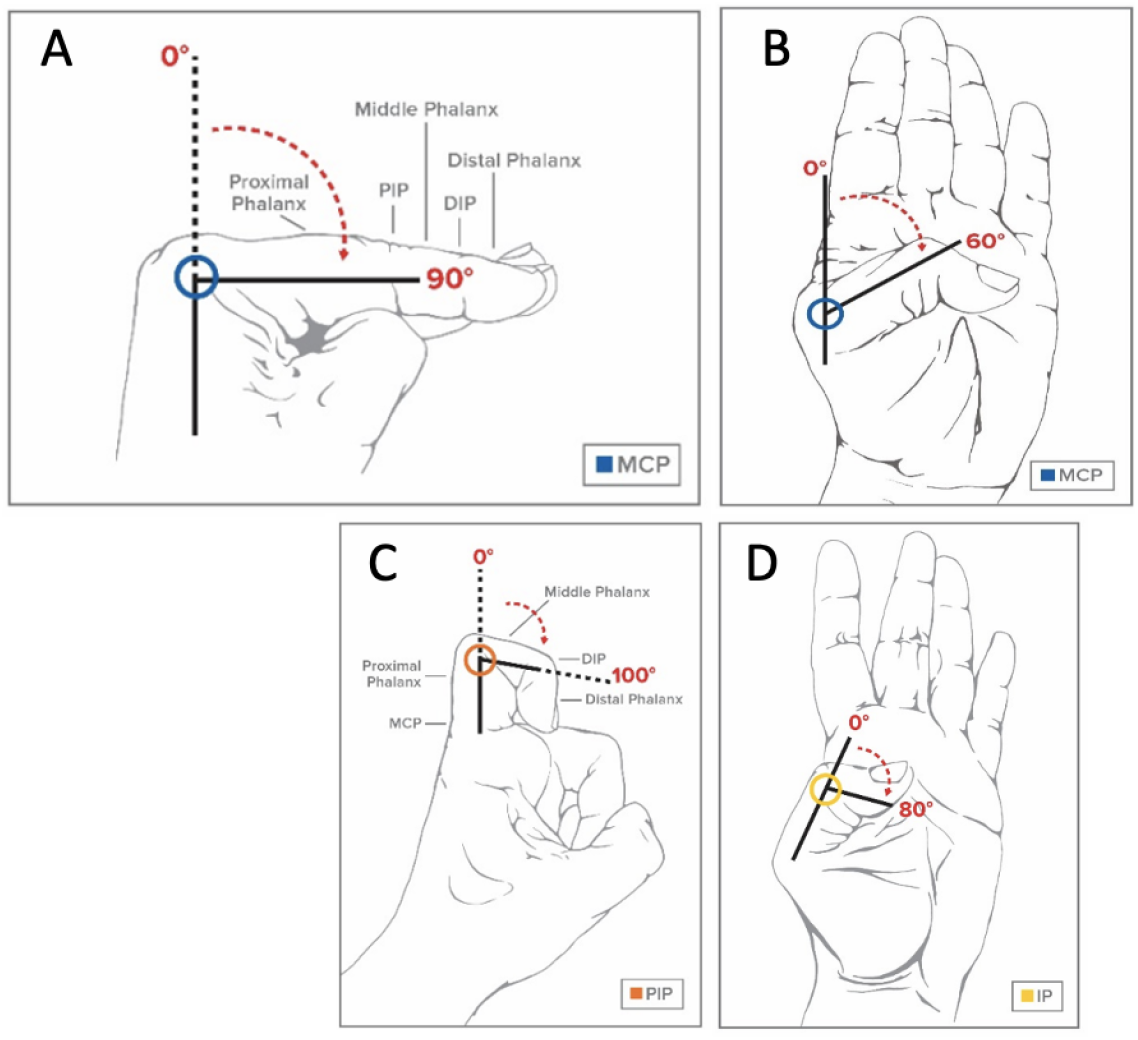
Measured angles. Measured position for the metacarpophalangeal (MCP) joint of the index finger (A), and of the thumb (B). Measured angles of the proximal interphalangeal (PIP) joint of the index finger (C), and of the thumb (D).

To describe the angles of the metacarpophalangeal joint, proximal interphalangeal joint, and distal interphalangeal joint, joints, the included angles between the segments were determined. Using the segments illustrated in Fig 11, the angles were calculated as:

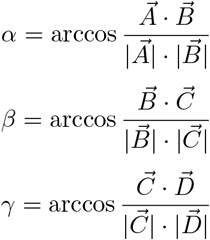

**Fig 11.**
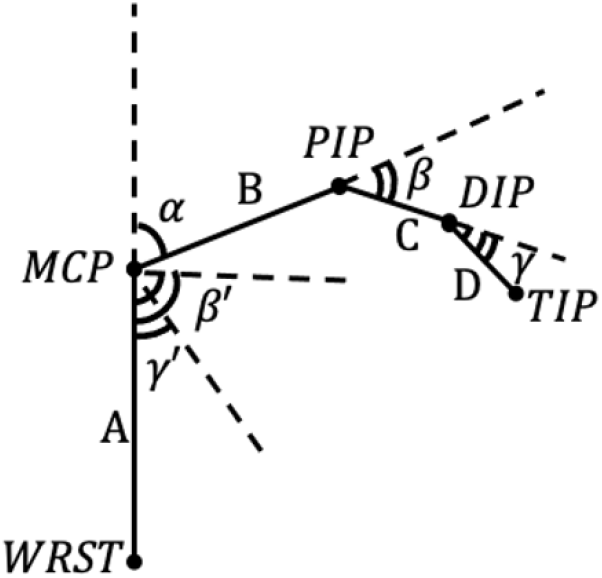
Geometric representation of the finger. Illustration of a geometric representation of the finger, where WRST indicates the wrist, MCP indicates the metacarpophalangeal joint, PIP indicates the proximal interphalangeal joint, and DIP indicates the distal interphalangeal. *α* represents the included angle of the metacarpophalangeal joint, *β* represents the included angle of the proximal interphalangeal joints, and *γ* represents the included angle of the distal interphalangeal joints.

A measurement to assess hand kinematics is the Total Active Flexion (TAF). Marx et al. on the journal of Hand Surgery defined the TAF as the measurement of active flexion of one digit [21]. Thus, TAF isolates the maximum flexion angle minus the minimum flexion angle, for a given activity, for metacarpophalangeal, the proximal interphalangeal joints, and the distal interphalangeal joints. Therefore, assessing the active flexion measures of joints under inspection for the specific exercise was selected as the preferred choice for this investigation.

## Acknowledgments

The authors confirm contribution to the paper as follows: study conception and design: LG, AB, AK; data collection: LG, WMRR; data analysis and interpretation of results: LG; draft manuscript preparation: LG, AB, AK.

## References

1. Field M, Pan Z, Stirling D, Naghdy F. Human motion capture sensors and analysis in robotics. Ind Robot Int J. 2011 Mar 8;38(2):163–71.

2. Metcalf CD, Notley SV, Chappell PH, Burridge JH, Yule VT. Validation and Application of a Computational Model for Wrist and Hand Movements Using Surface Markers. IEEE Trans Biomed Eng. 2008 Mar;55(3):1199–210.

3. Lopes TJA, Ferrari D, Ioannidis J, Simic M, Mícolis de Azevedo F, Pappas E. Reliability and validity of frontal plane kinematics of the trunk and lower extremity measured with 2-dimensional cameras during athletic tasks: A systematic review with meta-analysis. J Orthop Sports Phys Ther. 2018;48(10):812–22.

4. Reinking MF, Dugan L, Ripple N, Schleper K, Scholz H, Spadino J. Reliability of two-dimensional video-based running gait analysis.. Int J Sports Phys Ther. 2018;13(3):453.

5. Cao Z, Hidalgo G, Simon T, Wei SE, Sheikh Y. OpenPose: Realtime Multi-Person 2D Pose Estimation Using Part Affinity Fields.. IEEE Trans Pattern Anal Mach Intell. 2021 Jan 1;43(1):172–86.

6. Mathis A, Mamidanna P, Cury KM, Abe T, Murthy VN, Mathis MW. DeepLabCut: markerless pose estimation of user-defined body parts with deep learning.. Nat Neurosci. 2018;21(9):1281–9.

7. Macionis V. Reliability of the standard goniometry and diagrammatic recording of finger joint angles: a comparative study with healthy subjects and non-professional raters.. BMC Musculoskelet Disord. 2013 Jan 9;14(1):17.

8. Seethapathi N, Wang S, Saluja R, Blohm G, Kording KP. Movement science needs different pose tracking algorithms.. ArXiv Prepr ArXiv190710226. 2019.

9. Nakano N, Sakura T, Ueda K, Omura L, Kimura A, Iino Y. Evaluation of 3D markerless motion capture accuracy using OpenPose with multiple video cameras.. Front Sports Act Living. 2020;2:50.

10. Miller NR, Shapiro R, McLaughlin TM. A technique for obtaining spatial kinematic parameters of segments of biomechanical systems from cinematographic data.. J Biomech. 1980;13(7):535–47.

11. D’Antonio E, Taborri J, Palermo E, Rossi S, Patanè F. A markerless system for gait analysis based on OpenPose library.. In: 2020 IEEE International Instrumentation and Measurement Technology Conference (I2MTC). 2020. p. 1–6.

12. Sakurai T, Okada H. Examination of an applicable range for a markerless motion capture system in gait analysis. ISBS Proc Arch. 2021;39(1):141.

13. Stenum J, Rossi C, Roemmich RT. Two-dimensional video-based analysis of human gait using pose estimation.. PLoS Comput Biol. 2021 Apr 23;17(4):e1008935.

14. Drazan JF, Phillips WT, Seethapathi N, Hullfish TJ, Baxter JR. Moving outside the lab: markerless motion capture accurately quantifies sagittal plane kinematics during the vertical jump.. J Biomech. 125 (2021): 110547.

15. Sandau M, Koblauch H, Moeslund TB, Aanæs H, Alkjær T, Simonsen EB. Markerless motion capture can provide reliable 3D gait kinematics in the sagittal and frontal plane.. Med Eng Phys. 2014;36(9):1168–75.

16. Metcalf CD, Robinson R, Malpass AJ, Bogle TP, Dell TA, Harris C. Markerless motion capture and measurement of hand kinematics: validation and application to home-based upper limb rehabilitation.. IEEE Trans Biomed Eng. 2013 Aug;60(8):2184–92.

17. Li Z, Zhang R, Lee CH, Lee YC. An evaluation of posture recognition based on intelligent rapid entire body assessment system for determining musculoskeletal disorders.. Sensors. 2020;20(16):4414.

18. Pratt AL, Ball C. What are we measuring? A critique of range of motion methods currently in use for Dupuytren’s disease and recommendations for practice.. BMC Musculoskelet Disord. 2016 Jan 13;17:20.

19. Giavarina D. Understanding bland altman analysis.. Biochem Medica. 2015;25(2):141–51.

20. Metcalf CD, Robinson R, Malpass AJ, Bogle TP, Dell TA, Harris C, Demain SH. Markerless motion capture and measurement of hand kinematics: validation and application to home-based upper limb rehabilitation.. IEEE Transactions on Biomedical Engineering. 2013 Mar 7;60(8):2184–92.

21. Winter DA, Sidwall HG, Hobson DA. Measurement and reduction of noise in kinematics of locomotion.. Journal of biomechanics. 1974 Mar 1;7(2):157–9.

22. Marx RG, Bombardier C, Wright JG. What do we know about the reliability and validity of physical examination tests used to examine the upper extremity?. The Journal of Hand Surgery. 1999 Jan;24(1):185–93.

